# A scheme for 3-dimensional morphological reconstruction and force inference in the early *C. elegans* embryo

**DOI:** 10.1101/175166

**Authors:** Muzhi Xu, Yicong Wu, Hari Shroff, Min Wu, Madhav Mani

**Affiliations:** Engineering Sciences and Applied Mathematics, Northwestern University, Evanston, Illinois 60208, USA; Section on High Resolution Optical Imaging, NIBIB, NIH, Bethesda, Maryland 20892, USA; Mathematical Sciences, Worcester Polytechnic Institute, Worcester, Massachusetts 01609, USA; Molecular Biosciences, Northwestern University, Evanston, Illinois 60208, USA

**Author notes:** Equal corresponding authorship: M.W., M.M.

## Abstract

In this study, we present novel schemes for the reconstruction of cellular morphology and the inference of forces in the early *C. elegans* embryo. We have developed and bench-marked a morphological reconstruction scheme that transforms live-imaging of cellular membranes into a point cloud of smoothed surface patches, which facilitates accurate estimation of membrane curvatures and the angles between membranes.

Assuming an isotropic and homogeneous distribution of tensions along a membrane, we infer a pattern of forces that are 7% deviated from force balance at edges, and 10% deviated from the Young-Laplace relation at membrane faces. We have also demonstrated the stability of our scheme by sensitivity analysis of the coefficient matrices involved and the reproducibility of our image-analysis and force inference pipeline.

## Introduction

The emergence of morphology during organismal development, morphogenesis, consists of an interplay between biochemical signaling and mechanical forces. Despite our acquisition of an ever growing list of participating molecular factors, the collective nature of morphogenetic processes precludes a straightforward genotype-to-phenotype map. Furthermore, an abundance of *in vitro* studies have reported on the role of mechanotransduction in guiding cellular differentiation, suggesting that the map between chemical factors and physical forces is bidirectional. Understanding details of this map will constitute an advance in our conceptual understanding of morphogenesis.

Towards this end, single-molecule studies have provided a biophysical basis to the roles that biopolymers, adhesion molecules, and molecular motors play in morphogenesis. As such, morphogenetic processes involve the concerted and regulated action of a collective of adhesion and cytoskeletal proteins that prevents a straightforward understanding of the nature of forces given a list of participating molecules. The same molecules, in different regulatory states, can give rise to distinct morphogenetic phenomena. While central to the research agenda, the molecular approach faces difficulties in light of the complexity of regulation involved, and suggests taking a phenomenological and physical approach to morphogenesis.

Live fluorescence-based imaging facilitates pursuing a phenomenological approach to morphogenesis by giving us the ability to be quantitative about the geometry of cellular shapes and flows, and the dynamics of the cytoskeleton. However, given the complexity of a cell’s material properties, it is challenging to infer mechanical stresses from observed patterns of deformations and flows. In short, we lack the tools necessary to accurately and robustly measure the forces that determine cellular geometries and drive cellular flows in embryos. Recently, a set of image-analysis based indirect force inference schemes have begun to produce relative maps of forces in quasi two-dimensional epithelial tissues [1–7]. Force inference schemes are based on the assumption of force balance, are independent of the underlying material properties, and are constructed from the geometry of the tissue alone. The assumption of force balance is justified in settings where the timescales associated with cellular motion are large compared to the relaxation timescales observed following laser ablation events. The inferred cell-cell contact forces have been shown to correlate well with the average line density of molecular motor distributions [6]. It is worth noting that while the emergence of FRET-based reporters is exciting, connecting molecular-scale forces to the macroscopic forces that drive morphogenetic movements is a challenge [8].

The worm, *C. elegans*, embryo is an interesting setting within which to extend and apply image analysis based indirect force inference techniques. Juxtaposing *Drosophila* and *Xenpus* embryos, worm embryo’s undergo crucial cellular differentiation and morphogenetic processes with a small number of cells. Famously, the cell lineage of the worm is invariant, and is a consequence of both mosaic and regulative mechanisms. *What mechanisms impart the high degree of invariance in the worm’s cell lineage?* Manifestly, an answer to this question must take into account the biochemical and biophysical mechanisms at play in the embryo. Ironically, our biophysical understanding lags far behind our understanding of the biochemical dynamics alive in the worm.

Recent advances in the live-imaging of the *C. elegans* embryo gives unprecedented resolution to the complete 3D geometry and dynamics of the small number of cells as they make some of the most important and early decisions in the life of the worm. In this study, we present novel schemes for the reconstruction of cellular morphology and the inference of forces in the early *C. elegans* embryo. In particular, we present details of 1) an image analysis protocol that allows accurate reconstruction of the geometry of the membranes and junctions that facilitates 2) a scheme that gives access to the relative membrane tensions and cellular pressures over time in the *C. elegans* embryo. The enhanced accuracy of our morphological reconstruction was essential for inferring the desired tensions and pressures. Assuming an isotropic and homogeneous distribution of tensions along a membrane, we infer a pattern of forces that are 7% away from force balance at edges, and 10% away from the Young-Laplace relation at faces.

Furthermore, we present a sensitivity analysis that demonstrates the stability of our scheme. Lastly, we confirm that the reproducibility in the image-analysis pipeline is on the order of 5%. The quantitative assessment of the methodology presented in this study suggests improvements that we comment upon in our discussions section, and will guide future projects.

**Figure 1.**
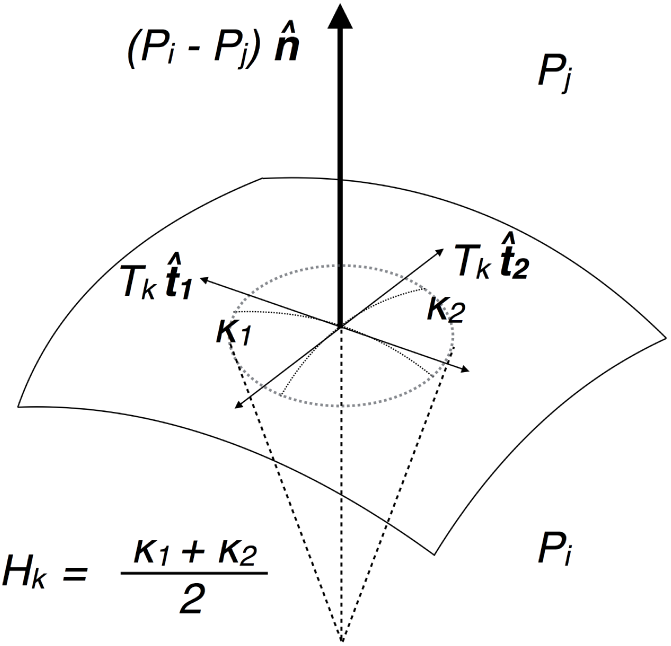
Schematic for Young-Laplace Force Balance on Membrane. The schematic depicts a small membrane patch, where normal pressure force on the membrane is balanced locally by the surface tension. Under isotropic tensions, the Young-Laplace relation, *P*_*i*_ *-P*_*j*_ = 2*H*_*k*_*T*_*k*_, is characterized completely by the mean curvature 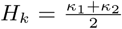. The two principle curvatures 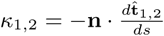, where **n** is the normal.

## Methods

### Force balance relations

We assume that the mechanical state of cells in the early worm embryo is dominated by intracellular pressures and intercellular membrane tensions. For each cell, indexed *c*, we define a pressure *P*_*c*_. For each membrane, indexed *m*, we define a tension *T*_*m*_. From a single frame from a movie of worm development we infer the unknown parameters (*P*_1_*,…, P*_*n*_*c, T*_1_*,…, T*_*n*_*m*) where *n*_*c*_ is the number of cells and *n*_*m*_ is the number of membranes. Furthermore, we ignore dissipative forces associated with the dynamics of the embryo underpinning our neglect of velocity data.

#### The Young-Laplace relation

We use the Young-Laplace relation on each membrane that relates the jump in pressure across a membrane to the product of its mean curvature and tension. The use of the Young-Laplace relation rests on the assumption that the membrane is fluid. We neglect inhomogeneities and anisotropies in tensions along a membrane, which is tantamount to assuming the variation of tensions along each membrane is negligible compared to the average tension. The following relation

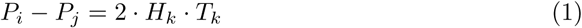

holds for each membrane face *k* (with adjacent cells *i* and *j*) where *P*_*i*_ and *P*_*j*_ are the pressures of cells *i* and *j*, respectively. *H*_*k*_ and *T*_*k*_ correspond to the mean curvature and the tension of face *k*, respectively (See Fig (1)). We have *n*_*m*_ Young-Laplace relations wherein the mean curvature *H*_*k*_ is obtained by taking the average of the mean curvatures of all points on the membrane. More details of calculating the mean curvatures are given in the *Image Analysis Protocol* section.

#### Junctional force balance

Curvilinear junctions are formed by the intersection between three membrane faces. At any point along these curvilinear junctions, we assume that the tension from the three intersecting membranes, should balance each other (See Fig (2)). Mathematically, the tensions *T*_*i*_, *T*_*j*_ and *T*_*k*_ are related by 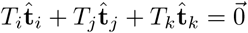 where 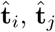 and 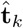 are the unit vectors normal to the junction, and tangent to the membrane surfaces *i*,*j*, and *k* respectively. The vector equation can be rewritten as

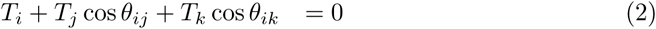

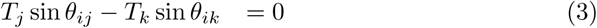

where *θ*_*ij*_ (*θ*_*ik*_) is the dihedral angle between 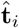 and 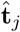 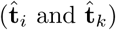, respectively. In principle, this relation holds at every point along the junction as the local tensions and angles vary along the junction. In this work, we define only one tension value for each membrane, so we only have one equation for each junction with constant *T*_*i*_, *T*_*j*_ and *T*_*k*_ values and *θ*_*ij*_ and *θ*_*ik*_ are taken as the averages along the junction.

**Figure 2.**
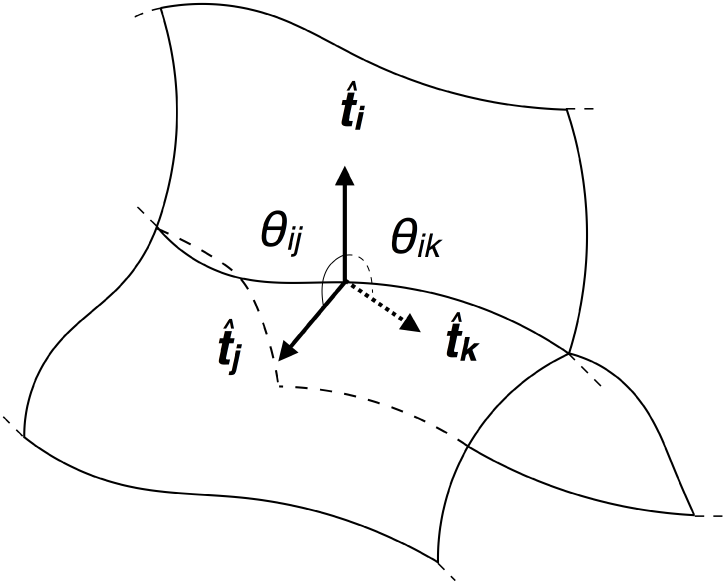
Schematic for Tension Balance at Edge Junction. The three membrane faces, illustrated as curved planes, intersect at the edge junction. The surface tensions act perpendicular to the edge junction under isotropic tensions. The dihedral angle of intersections between the faces prescribe the relation in Eq. (2) and (3).

#### Solving the system of equations

We define the vector

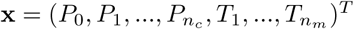 where *P*_0_ is the constant pressure value exterior to the embryo, 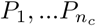 are the pressure values for the cells and 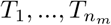 are the tension values for the membranes. We can solve the linear system

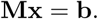

for the pressure and tension values where the size of **M** is

(*n*_*m*_ + 2 *× n*_*j*_ + 2) *×* (*n*_*m*_ + *n*_*c*_ + 1). The first *n*_*m*_ rows of **M** correspond to the force balance relations along the membrane by Eq. (1) and the next 2*×n*_*j*_ rows correspond to the force balances along the junctions by Eq. (2) and (3). The two additional rows come from two equations 1) fixing the exterior pressure, *P*_0_ = *P*_*b*_, and 2) setting the scale of tension values. In this work, we scale the average tension value to be 1 – the equation for setting the scale for the tension is 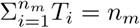. Finally, we have **b** = (0*,…,* 0*, P*_*b*_*, n*_*m*_)^*T*^. Notice the relation between the tension and pressure from the

Laplace’s Equation depends on the length scale of the image since they hold different units. In practice, we can set the tension scale and length scale, which effectively determines the pressure scale. Based on the inequality *n*_*m*_ + *n*_*c*_ + 1*≤n*_*m*_ + 2*×n*_*j*_ + 2 (can be proved by induction from a single cell, where 1 + 1 + 1 *≤* 1 + 2 *×* 0 + 2 holds), the system is overdetermined and we can only solve **x** in the sense of minimization of the error ∥**Mx** *-* **b** ∥ ^2^, which is equivalent to 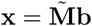 where 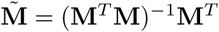 is the pseudoinverse of **M**.

### Imaging

Nematode strain BV24 ([ltIs44 [pie-1p-mCherry::PH(PLC1delta1) + unc-119(+)]; zuIs178 [(his-72 1kb::HIS-72::GFP); unc-119(+)] V]) was used for membrane imaging. ltIS44 is an integrated transgene which expresses membrane-localized mCherry; zuIS178 is an integrated transgene that expresses histone-GFP fusion, which was not imaged in this study. Worms were raised under standard conditions at 20^*?*^C on NGM media seeded with E. coli OP50. The embryo was transferred to rectangular coverslips (VWR, 48393-241) and then placed into an imaging chamber (Applied Scientific Instrumentation, I-3078-2450) as previously described1. The embryo was imaged with dual-view inverted selective-plane illumination microscopy (diSPIM)2, although only one view was selected for analysis. The illumination wavelength was 561 nm (Crystalaser, CL-561-050), mCherry fluorescence was collected via the 0.8 NA detection objective (Nikon 40*×*, 3.5 mm working distance, water immersion lens) transmitted through dichroic mirrors (Chroma, ZT405/488/561rpc), filtered through a notch emission filters (Semrock, NF03-561E-25) to reject the 561-nm pump light, and imaged with 200-mm tube lenses (Applied Scientific Instrumentation, C60-TUBE B) onto scientific-grade, complementary, metal-oxide semiconductor (sCMOS) cameras (PCO, Edge). The resulting image pixel size was 6.5 *μ*m/40 = 162.5 nm. We recorded 80 planes per volume for the embryo, 5 ms per plane, spacing planes every 0.5 *μ*m. Volumes were recorded at a temporal resolution of 1 min from 4-cell stage until hatching (i.e., 13 hours post fertilization).

### Morphological reconstruction of the embryo

The coefficient matrix **M** involves the averaged mean curvatures *H*_*k*_ over each k-th membrane surface and the averaged dihedral angle *θ*’s between intersecting membranes along the curvilinear junctions. Before we compute the averaged curvatures and angles, we need to reconstruct these parameters locally at each membrane and junction point. But these parameters, especially the curvatures *κ*_*m*_, which involve the second-order derivatives of position vectors along the membrane surfaces, are sensitive to noise.

Noise can arise during the creation of the gray-scale image due to noise inherent to fluorescence imaging. Instead of working with the gray-scale image, we work with the probability map generated by the pixel classification scheme of the machine learning image analysis software (*Ilastik*). We show two samples of the grayscale image and the probability map in Figure 3 (A, B and E), where the first two rows in panel A, E and B show the the cross-sectional image of the 7-cell-embryo and 12-cell-embryo, together with 3D image of the 7-cell-embryo, respectively. The third rows in panel A and E show a cross section of the segmented one-voxel-thick membrane structure. We can observe that the reconstructed membranes are artificially distorted or flattened due to their restriction on the voxel mesh. The distortion can result in an unreliable estimate of the normal vectors and in an inaccurate computation of the curvatures along the membrane’s surface. Moreover, the junctions reconstructed by taking the intersection of three adjacent membrane surfaces are also distorted and can disrupt the computation of the tangent vectors of the junctions and the angles between intersecting membranes along junctions. See the first row of panel D in Figure 3 of the disorganized tangent vectors.

To deal with this artifact, we smooth the membrane surfaces based on a principal component analysis (PCA) of the membrane points obtained by the watershed transformation. The smoothed membrane surfaces are reconstructed on a three-dimensional point cloud, a collection of mesh-free points. The outcome of this protocol can be seen in the fourth row of panel A and the second row of panel D from Figure 3, compared with their unsmoothed counterparts above. The details of this method are discussed in the section, *Point cloud normal estimates and smoothing*. Based on the smoothed membrane surfaces and junctions, we compute the curvatures and the angles between intersecting membranes, detailed in *The curvatures of the membrane surfaces* and *The angles between membranes along the junctions*, respectively.

**Figure 3.**
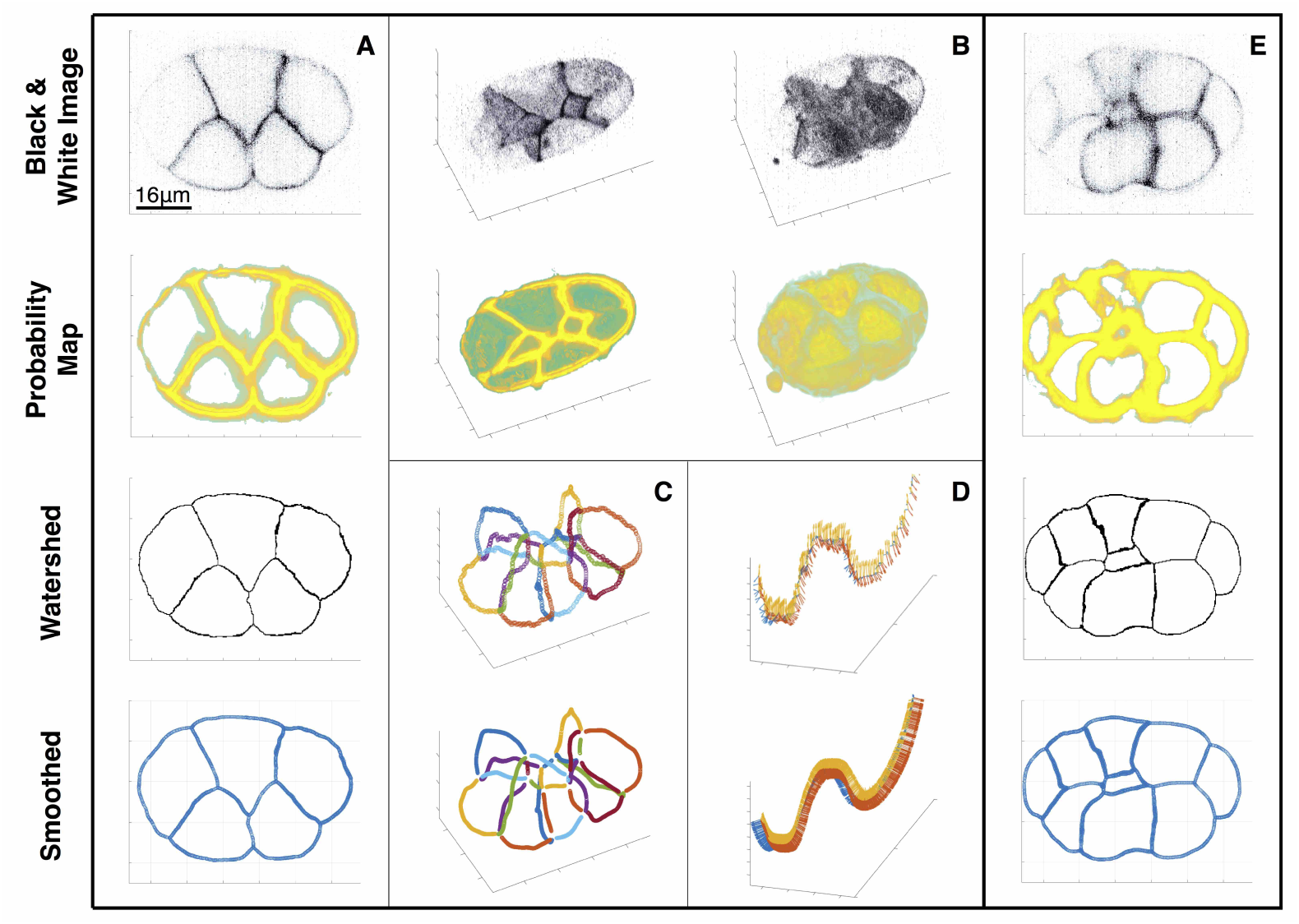
Workflow of Embryo Reconstruction. (The cross section is taken near the top where only 5 cells are shown. Figure 3 (Panel A and B in the first two rows) shows the cross-sectional and three-dimensional image of the a 7-cell-embryo sample of the grayscale image and the probability map. **A**) Progression of the a cross section from the middle of the embryo. In the probability map, the yellow and green indicate high and low probabilities respectively. (**B**) Visualization of the black & white image and probability map, with a cross section (left) and the whole embryo (right). (**C**) Plot of all the edge junctions in the embryo. The effect of smoothing can be seen clearly here. (**D**) Depiction of an edge junction along with the tangent vectors of the adjacent membrane faces. The sparser watershed edge junction is replaced by a denser and less noisy point cloud representation. (**E**) Plot of the edge junction located on the bottom of the embryo, with the face tangent vectors. Angles between faces remain mostly unchanged.

#### Point cloud normal estimates and smoothing

The surface normal vectors are required to compute the curvatures and the angles between the intersecting membrane surfaces. We estimate the surface normal by the method from [9, 10], which smooths the surface locally and estimates the normal simultaneously. Given that the surface is sufficiently smooth, the surface normal at a point **p** can be obtained by finding the unit vector ∥***η***∥ = 1, which minimizes 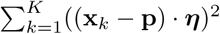, where **x**_*k*_’s are the *K*-nearest neighboring points of **p** (see *K*_*S*_ in the Parameter Table in SI). It is equivalent to the normal estimate method from principal component analysis (PCA). In order to smooth the surface, a shift *t* is introduced in [9, 10] along the unknown normal direction ***η***, and the constrained least square problem can be reformulated to: Find *t* and ***η*** that minimize

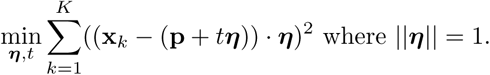

at the shifted 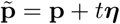. After computing the surface normal vectors at every point along the surface, there is no guarantee that their orientation will be consistent. We follow the procedure in [9–11] to propagate the consistent direction of the normal vectors along the Euclidean minimum spanning tree that connects the points.

#### Estimating membrane curvatures

We calculate the mean curvature *κ*_*m*_(**p**) at each membrane point **p** = (*x*_*p*_*, y*_*p*_*, z*_*p*_) by first parametrizing the membrane patch including the K-nearest neighboring points **x**_*k*_ = (*x*_*k*_*, y*_*k*_*, z*_*k*_)’s based on the normal estimate ***η*** and smoothing from the last section. At each point, we define a local *u -v -z* cartesian coordinate system where *u* and *v* span the tangent plane and *z* is along the normal direction ***η***. Then we parametrize the surface patch by **r**(*u, v*) = *u***t**_1_ + *v***t**_2_ + *z*(*u, v*)***η***, where **t**_*i*_’s are the unit tangent vectors. *z*(*u, v*) can be approximated by fitting the the local surface by a second order Taylor expansion about **p**, minimizing

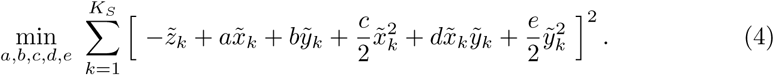

where 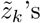 are the projections of (**x**_*k*_ *-* **x**_*p*_)’s along ***η***, and 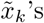 and 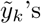 are their projections in the tangent plane. Then the curvatures can be computed from the shape operator *W*_*p*_, which is a 2 *×* 2 matrix (also called Weingarten matrix) where the components can be calculated by differentiating **r**(*u, v*) up to the second order. See more details in the SI. The local mean curvature *κ*_*m*_ is calculated by taking the average of the two eigenvalues of the matrix (principal curvatures). We also further update the normal 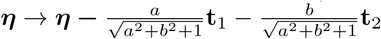 accordingly.

#### Estimating the angles between membranes

On the points along the membrane junctions, we define the local angles between the three intersecting membranes in the plane normal to the tangent direction (normal plane, see the blue plane in Figure 4). We need to 1) reconstruct the junction, 2) estimate the normal plane and 3) compute the angles.

To reconstruct the junction for each of the three intersecting membranes we sample the points into a temporary point cloud within a distance threshold *d*_*T*_ (see the parameter table in SI) from the other two membranes. On the temporary point cloud, we update the position of each point by the average of the position vectors of the K-nearest neighboring points (see *K*_*J*_ in the parameter table in SI). This update turns the temporary bold point cloud into a thinned junction curve.

To estimate the normal plane along the thinned junction curve, we define the tangent vector on each point along the junction curve approximated by

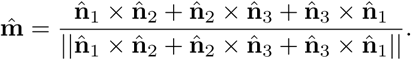

where 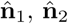, and 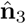 are the surface normals evaluated on the three closest points from the three membranes. We then define the adjusted normals 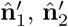, and 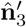 by

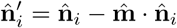

and the adjusted normals are in the normal plane, as shown in Figure 4. Based on the adjusted normals, we can compute surface tangents 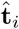 orthogonal to the junction curve by

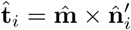

for membrane face *i*. The angles *θ*_*ij*_ and *θ*_*ik*_ from Eq. (2) and (3) are finally computed by taking the differences between 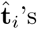.

**Figure 4.**
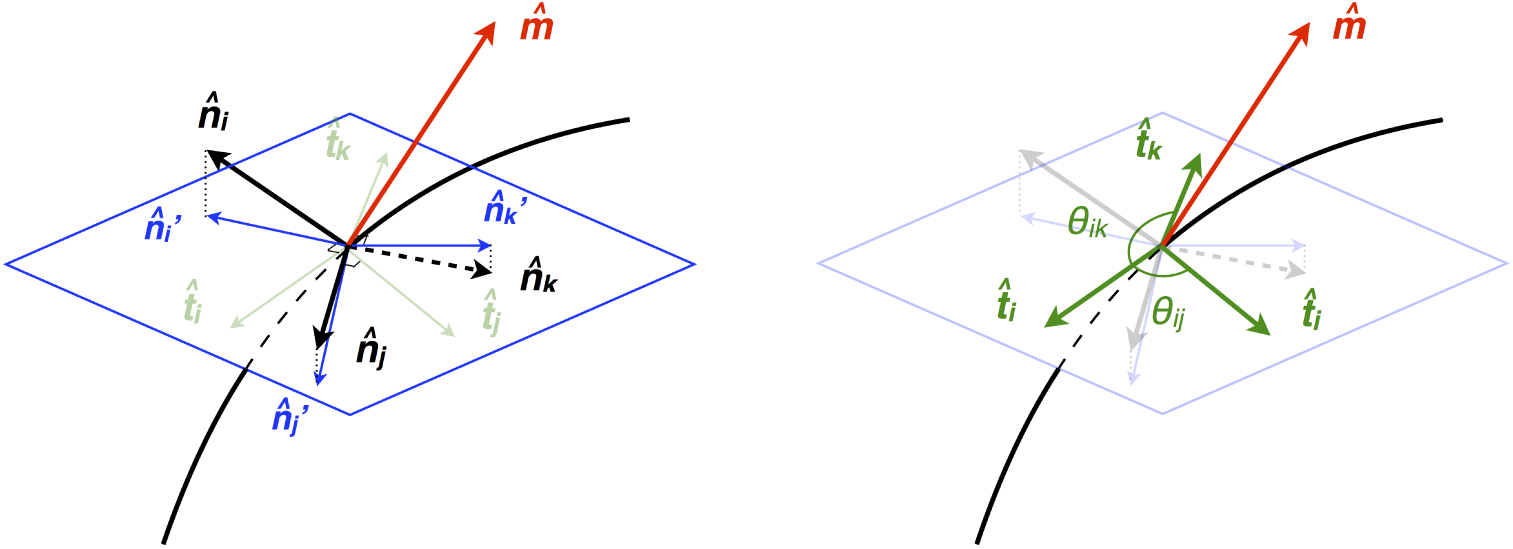
Schematic for Junction Tangent and Angle Computation. The black curved line represents the edge junction and the black arrows tagged by 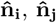, and 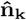 represent the normal vector of each membrane face. The red vector 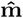 is the approximation to the edge tangent vector. The blue plane represents the perpendicular plane to the red vector 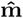. The blue vectors are the projected normals which lie in the plane normal to 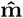 and thus are coplanar. On the right, it is shown that the angle of intersections can be calculated from the tangent vectors 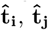, and 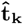 from each membrane face normal to the edge junction.

## Results

### *In-silico* validation of the scheme

Our model assumes that the mechanical state of cells in the early worm embryo is dominated by intracellular pressures and intercellular membrane tensions, which are isotropic and uniform on each membrane face. Before we implement the workflow to the worm embryo, we test it against an *in-silico* two-cell systems where the above assumptions are fully satisfied. In detail, the configuration of a two-cell system can be fully determined by the tensions from the three membrane faces and pressures from the two cells. We generate a family of synthetic membrane images where the radii of the two cells (*R*_1_ = 5 *× L* and *R*_2_ = 4 *× L*) and the radius of closed circular junction (*d* = 3 *× L*) are fixed (*L* is the length scale). See the left panel in Figure 5 for the schematics. By changing the ratio of pressures (*P*_2_*/P*_1_) between the two cells and the ratio between tensions accordingly, we can maintain the radii of the two cells while changing the mean curvature of the interfacial membrane (*H*_3_). For this family of configurations with different *P*_2_*/P*_1_’s, we synthesize 3D images with probability maps of membrane faces with different spatial resolutions. We choose *L* = 10, 20, 40 to effectively change the resolutions. *L* = 10 corresponds to approximately 45 voxel numbers across the cell diameter while *L* = 40 corresponds to approximately 182 voxel numbers across the cell diameter. Based on the probability maps, we reconstruct the cell membranes into a point cloud and extract the curvatures of the three membranes 1*/R*_1_, 1*/R*_2_ and *H*_3_ and the angles *θ*_12_,*θ*_13_ and *θ*_23_ between the three membranes along the circular junction. The reconstruction of the curvatures of the two major membranes 1*/R*_1_ and 1*/R*_2_ are reliable regardless of the different resolutions used (see errors between the reconstructed and true curvature in Fig S1,A), while the curvature reconstruction of the interfacial membrane *H*_3_ is improved by increasing the resolution (see Fig S1,B). Notice as *P*_2_*/P*_1_ approximates to 1, the curvature of the interfacial membrane *H*_3_ approaches to 0, and this is why the relative errors elevate as *P*_2_*/P*_1_ decreases in Figure 5B. The reconstruction of angles *θ*_12_,*θ*_13_ and *θ*_23_ is also improved by increasing *L* (see Figure S1,C for the total error 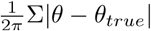). With respect to our force inference scheme, we show that the total relative error 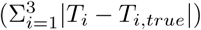 between the measured tensions and the true tensions is below 0.2 in all cases and decreases when the resolution increases (See the right panel in Figure 5). We also show that the relative residuals of equations from the Young-Laplace relation and the force balance along the junction (normalized as described in the results *Quantitative assessment of errors*) are below 0.02 and 2 *×* 10^−5^, respectively (See Figure S1,E and F).

**Figure 5.**
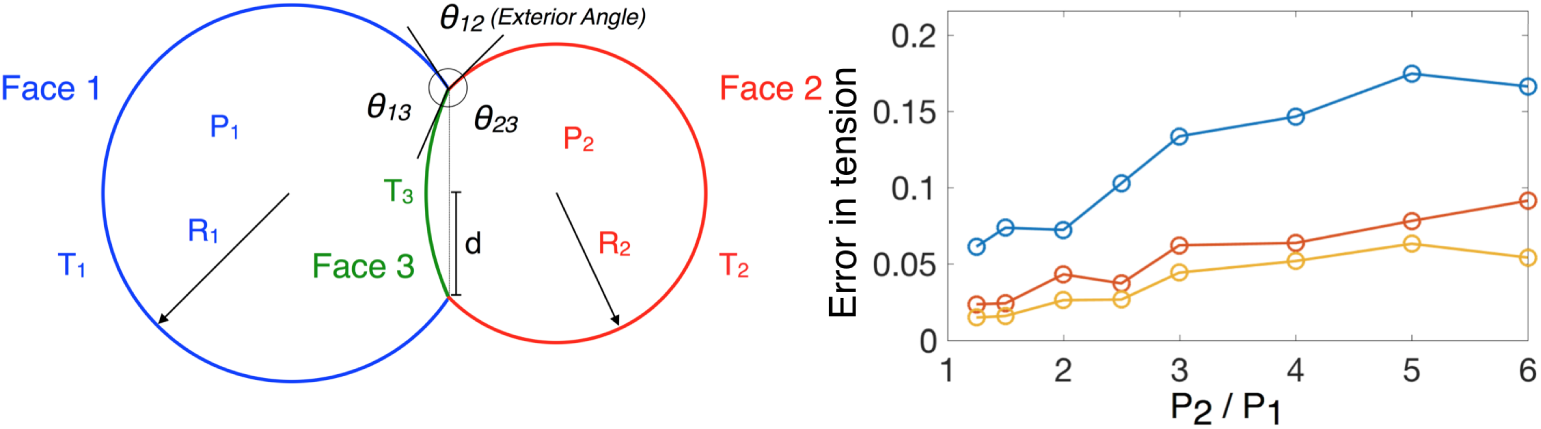
Schematics of a two-cell system and error in tension inferences. On the left, the system depicts the cross-sectional slice of two cells (with pressures *P*_1_ and *P*_2_) with constant mean curvatures *H*_1_ = 1*/R*_1_ and *H*_2_ = 1*/R*_2_ on the major membrane faces 1 and 2 (with tensions *T*_1_ and *T*_2_) separated by the interfacial membrane face 3 (with tension *T*_3_) with constant mean curvature *H*_3_. Note that all three membranes are patches of spherical membranes as they have constant mean curvatures. *d* denotes the radius of the circular junction between the 3 membrane faces and the *θ*’s denote the angles between the 3 membranes along the circular junction. On the right, we measure the total error between the true tension and the inferred tension: 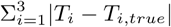.

### Morphological reconstruction of worm embryos

See Figure 6 for the reconstructed mean curvatures *H*_*k*_’s of the membrane faces and averaged dihedral angles along the curvilinear junctions *θ*’s using the workflow *Morphological reconstruction of the embryo*. We obtain the average mean curvature *H*_*k*_ on each membrane face *k* by taking the average of local mean curvatures, *κ*_*m*_, on each membrane face. See Figure 6 (A) for the heat map of *κ*_*m*_ on an exterior membrane face. Since the local mean curvature at a point **p** is obtained by fitting the neighboring membrane points by a paraboloid (see Eq. 4 in *The curvatures of the membrane surfaces*), the result changes with *K*_*C*_, the number of nearest neighboring points used. However, the averaged mean curvature *H*_*k*_ is not sensitive to the choice of *K*_*C*_. See Figure 6 (B) for the distributions of mean curvatures *κ*_*m*_ using *K*_*C*_ = 50, 800 and 3200. Taking from here, we fix *K*_*C*_ = 50 for each membrane to calculate the *κ*_*m*_’s and the associated *H*_*k*_’s. See Figure 6 (D-F) and 6 (G-I) for the averaged mean curvature *H*_*k*_’s on the exterior membranes and interior membranes in different views. See Movie 1 and 2 for more details of *H*_*k*_’s in both the 7-cell-embryo and 12-cell-embryo. Note that the curvature we obtain is in units of voxel sizes, and later we directly use the non-dimensional curvature to infer relative pressures. Similarly, we calculate the averaged dihedral angles between membranes along the curvilinear junction from the local dihedral angle computations following *The angles between membranes along the curvilinear junctions*. See Figure 6 (C) for the angle variations along one junction intersected between two exterior membranes and one interior membrane. With respect to our proposed force inference scheme we obtain all the parameters needed in the coefficient matrix **M** to infer forces in both 7-cell-embryo and 12-cell-embryo.

**Figure 6.**
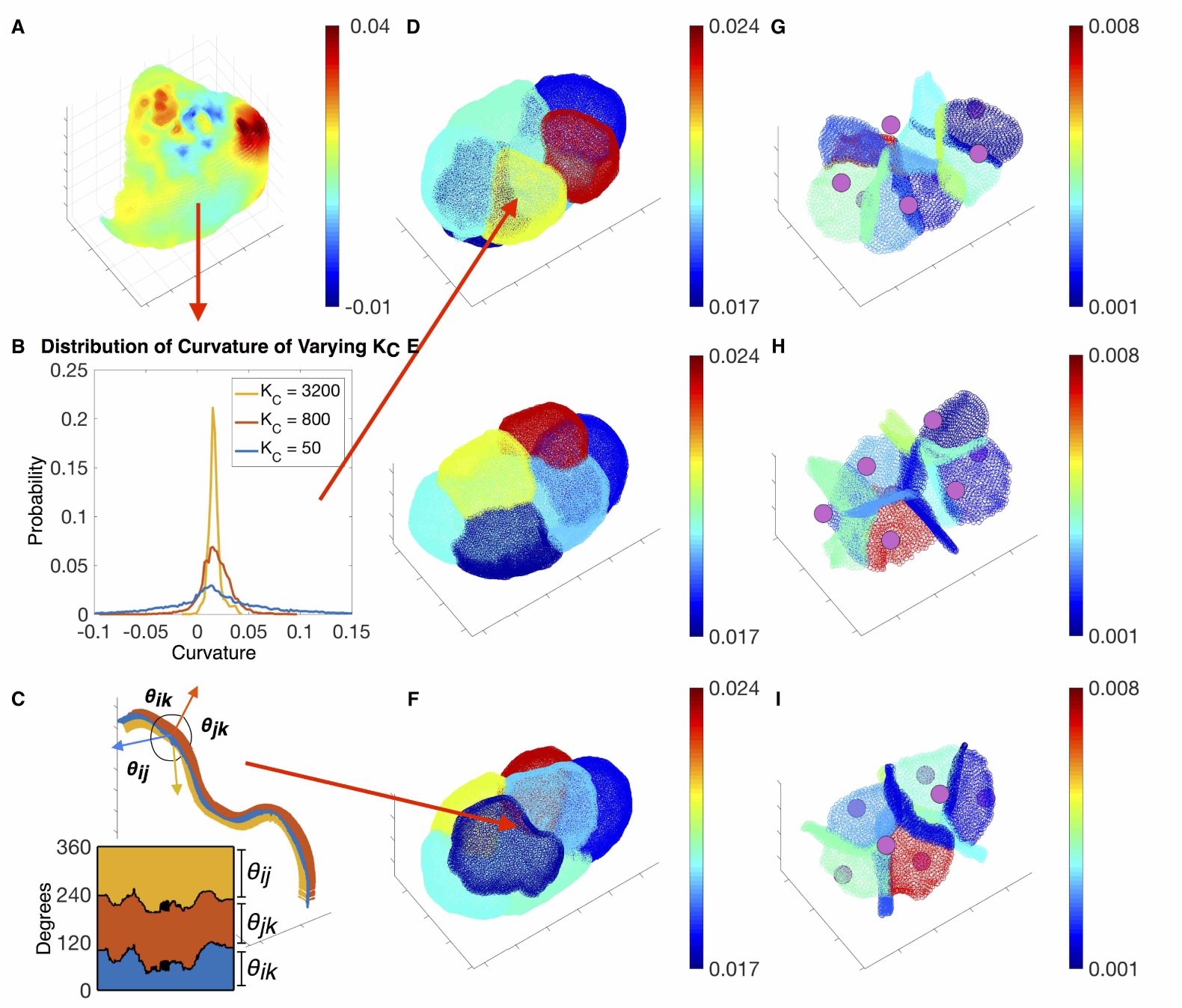
The reconstruction of curvatures and angles. (D-F) show the average mean curvatures *H*_*k*_’s on the exterior membranes and (G-I) show the average mean curvatures on the interior membranes by rotating the anterior-posterior axis in different angles. See movie 1 and 2 for the average mean curvatures for both 7-cell-embryo and 12-cell-embryo. The average mean curvature *H*_*k*_’s on each membrane face is calculated by taking the average of mean curvatures *κ*_*m*_(**p**) over all points on the same membrane face. A shows an example of mean curvature distributed on a single membrane face and B shows the distribution of mean curvatures changes by using different number of nearest neighboring points *K*_*C*_, but the overall averaged mean curvature is not sensitive to the choice of *K*_*C*_. From here, we always use *KC* = 50 (also see the parameter table) to calculate the average mean curvatures *H*_*k*_’s on each membrane face. We calculate the three average angles *θ*’s along each junctions by taking the average of three angles over all points on the same junction. C shows an example of the angle variations along one junction.

**Figure 7.**
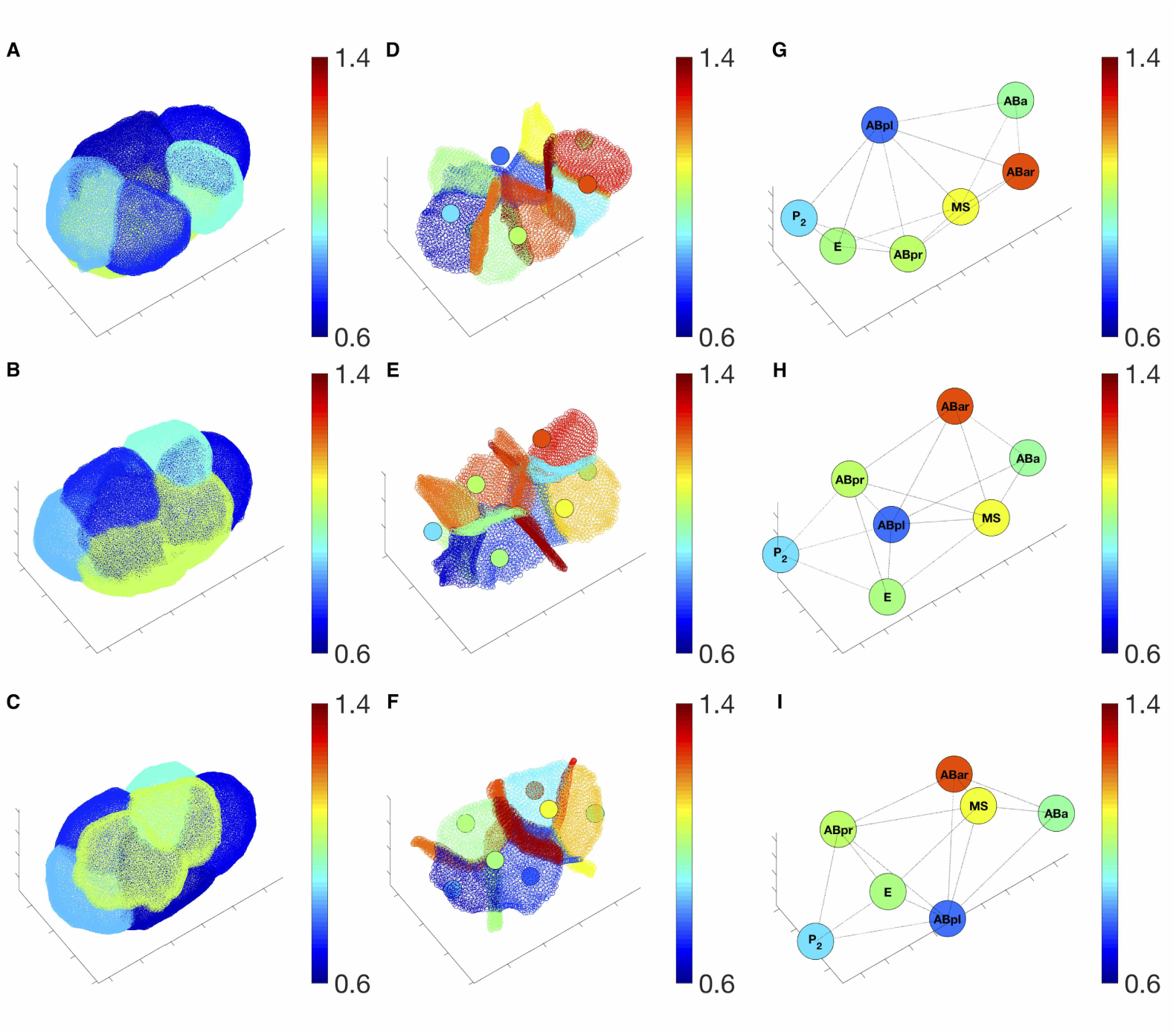
Inferred Forces. Depiction of the inferred tensions and pressures at the 7 cell stage. (A-C) depict relative tensions on the outer membranes. (D-F) depict the relative tensions on the inner membrane and the relative pressures in the cells (represented as colored circular region in the middle of each cell). (G-I) depict the pressures in the cells labeled by their names. For presentation, all pressure values are rescaled to match the same scale as the relative tension. The three rows depict the embryo viewed at different angles, where the embryo is rotated around an axis along its anterior to the posterior.

### Force inference on the embryo

We infer the average tensions on the membrane faces and pressures in cells according to the model in *Force balance relations*. The pressure in each cell and the average tensions on each membrane are shown in Figure 7 from three different views. The pressures are shown both in the second column (panel D-F) together with the tension map of the inner membranes as well as in the third column (panel G-I) where each cell is labeled by its name. The ABar cell on the anterior of the embryo sustains the highest level of pressure while the ABpl cell on the posterior sustain the lowest level of pressure.

Interestingly, the P2 cell, which is about to divide at this stage in the movie, shows a relatively low level of pressure. A global monotonic gradient of pressure from the right anterior side to the left posterior side of the entire embryo can be discerned. In contrast, we cannot identify a global transition of tensions among the inner membranes from the anterior to the posterior side of the embryo, and the distribution of tensions on the inner membranes are heterogeneous. However, one can see that most of the inner membranes (panel D-F) present higher level of tensions than the outer membranes (panel A-C). We conjecture that this is likely a consequence of outer membranes only comprising approximately half the cytoskeleton activity that inner membranes posses. See Movie 3 and 4 for more details of inferred forces in both 7-cell-embryo and 12-cell-embryo.

Based on the inferred tension and pressure, we can further reconstruct the averaged mechanical state (the full stress tensor) of each cell according to

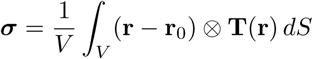

where *V* and **r**_0_ are the volume and the centroid of the cell, respectively, and **T**(**r**) is the traction at location **r**. See Figure 8. Physically speaking, the cell’s stress tensor,***σ***, is obtained by integrating the traction **T**(**r**), force per area, acting on every point, **r**, on the cell membranes averaged by volume. The three orthogonal principal axes of the stress tensor point in the directions that the cell is shear-free, and the eigenvalues quantify the uniaxial tension along each principle direction. In Figure 8, we plot the principle axes of stress tensors on each cell, where the lengths of the axes show the relative magnitude of the tensile stress in the corresponding direction. One can see that the ABpl and ABpr cells are exposed to high levels of stress and stress anisotropy. See Movie 5 and 6 for more details of the stresses in both 7-cell-embryo and 12-cell-embryo. Similarly we can also reconstruct the shape tensor of each cell to quantify its relative size compared to other cells and its shape anisotropy. The shape tensor is computed by rescaling the moment of inertia tensor by volume:

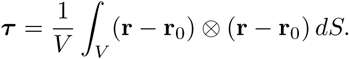

The three orthogonal principal axes of the shape tensor resemble the principal axes of an ellipsoid and the ratios between the eigenvalues quantify the shape anisotropy of the cell. In summary, here we have inferred forces in both 7-cell-embryo and 12-cell-embryo and have visualized their mechanical state accordingly.

### Quantitative assessment of errors

Here we present a sensitivity and reproducibility analysis of the proposed schemes. Errors in the inferred forces from the equilibrium solution arise from four main sources. First, noise can be introduced due to photon noise during image acquisition. Second, noise can be introduced via our segmentation protocol. Third, errors might be accrued by our modeling assumption were they to inaccurately represent the mechanical state of the embryo. Fourth, our method solves an overdetermined system, and as such not every balance relation can be fully satisfied. Below, we will first assess and discuss the errors of our inferred forces for the balance relations due to the overdetermined system. Then in *Sensitivity analysis and reproducibility of protocol*, we will discuss the robustness of our method to noise from the first two steps. The error due to the assumption of our model can be tested via a correlation study of the average myosin distributions along each membrane with our predicted membrane tension, which will be conducted in a future study.

We quantify the relative errors on the membranes and junctions separately. On each membrane, the absolute error is defined as the residual of the Young-Laplace relation (1). The relative error is obtained by (*P*_*i*_ − *P*_*j*_ − 2 *· H*_*k*_ *· T*_*k*_)*/*(*|P*_*i*_ *|*+ *|P*_*j*_*|* +*|* 2 *· H*_*k*_ *· T*_*k*_*|*) -the residual of equation (1) divided by the magnitude sum of each term. A scatter-plot of the left and right hand side of equation (1) is reproduced in Figure 9 (A). The two clusters in the scatter plot correspond to the force balance equations on the outer (Figure 9 (B)) and inner membranes (Figure 9 (C)). The clustering can be explained by the fact that the outer membranes have higher mean curvatures on average than the inner membranes. The colors of the points visualize the magnitude of the relative errors, which is also plotted in the same color code on the outer and inner membranes below. We note that the largest errors are concentrated on the anterior outer and inner membranes. The average relative error for the outer membranes is 11.2% and for the inner membranes is 9.12%.

On each junction, the force balance is described by equations (2) and (3) in the two orthogonal directions. We define the absolute force balance error by the magnitude of the residual vector from the two equations. The relative error is then obtained by rescaling the absolute error by the average total forces among all the junctions, where the total force on each junction is the summation of the three inferred tensions. We plot the errors among all the junctions both in the histogram (Figure 9 (D)) and in the heat map (Figure 9 (E)). We note that the largest error occurs on the shortest outer junction. The average relative error over the junctions is 7.21%.

### Sensitivity analysis and reproducibility of protocol

Here we perform sensitivity analysis to estimate the reliability of the results subject to noise. Under small perturbation of the coefficient matrix **M** + *δ***M** and **b** + *δ***b**, we can look at the spectrum (i.e., the set of eigenvalues *λ*_*i*_’s, *i* = 1, 2*,…, n*_*m*_ + *n*_*c*_ + 1) of the pseudoinverse 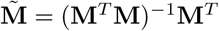 to estimate the sensitivity of **x** + *δ***x** to *δ***M** and *δ***b**. Large *λ*_*i*_ *>* 1 indicates *δ***x** is sensitive to the perturbations. We perform the sensitivity analysis to the 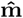 of a 7-cell embryo and a 12-cell embryo and both of them show most of the eigenvalues are smaller than 1. (See the spectrum distribution in Figure 10) Interestingly, the eigenvector corresponding to the largest eigenvalue is in the direction of constant value for all pressures and zero for all tensions. Since we are looking at the pressure difference from the exterior pressure, this perturbation mode does not contaminate the result. To check the reproducibility of our workflow, we have reconstructed a grayscale image of the membranes from the smoothed point cloud based on which we calculate **M** by 1) generating a black-and-white image by rounding-off the positions of the point cloud to the nearest voxels and 2) diffuse the membrane voxels 3-voxel-distance away followed by a linear decrease of the intensity value. This effectively generates an image with the same intensity profile away from the membrane points as the original data. We run our workflow on the regenerated data and find the relative error between the result from the original and the regenerated data for the 7-cell-embryo below 5%. See the inferred pressure and tension using reprocessed data in Figure 10 vs forces using the original data.

**Figure 8.**
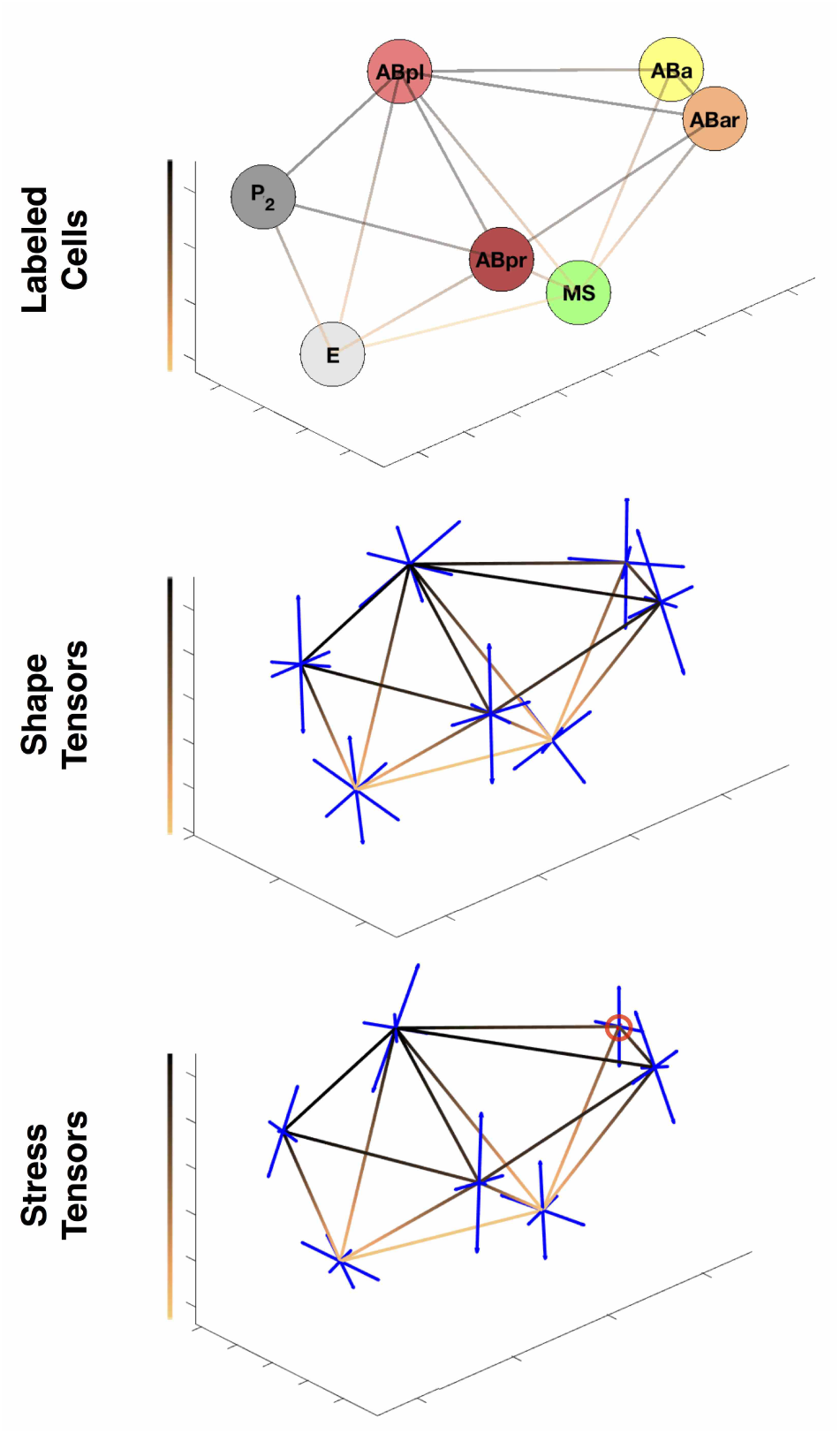
Depiction of average mechanical state and shape tensors. Plot of the shape and stress tensors at the 7 cell stage. The copper gradient lines represent cell connectivity, while the color ranging from black to copper corresponds the depth in z-axis. The tensors (all of which are symmetric) are represented by their 3 orthogonal eigenvectors plotted as blue line segments. The length of the segments correspond to the magnitude of the eigenvalue. Compressive forces in the stress tensors are plotted as red lines, and are circled in red for clarity.

**Figure 9.**
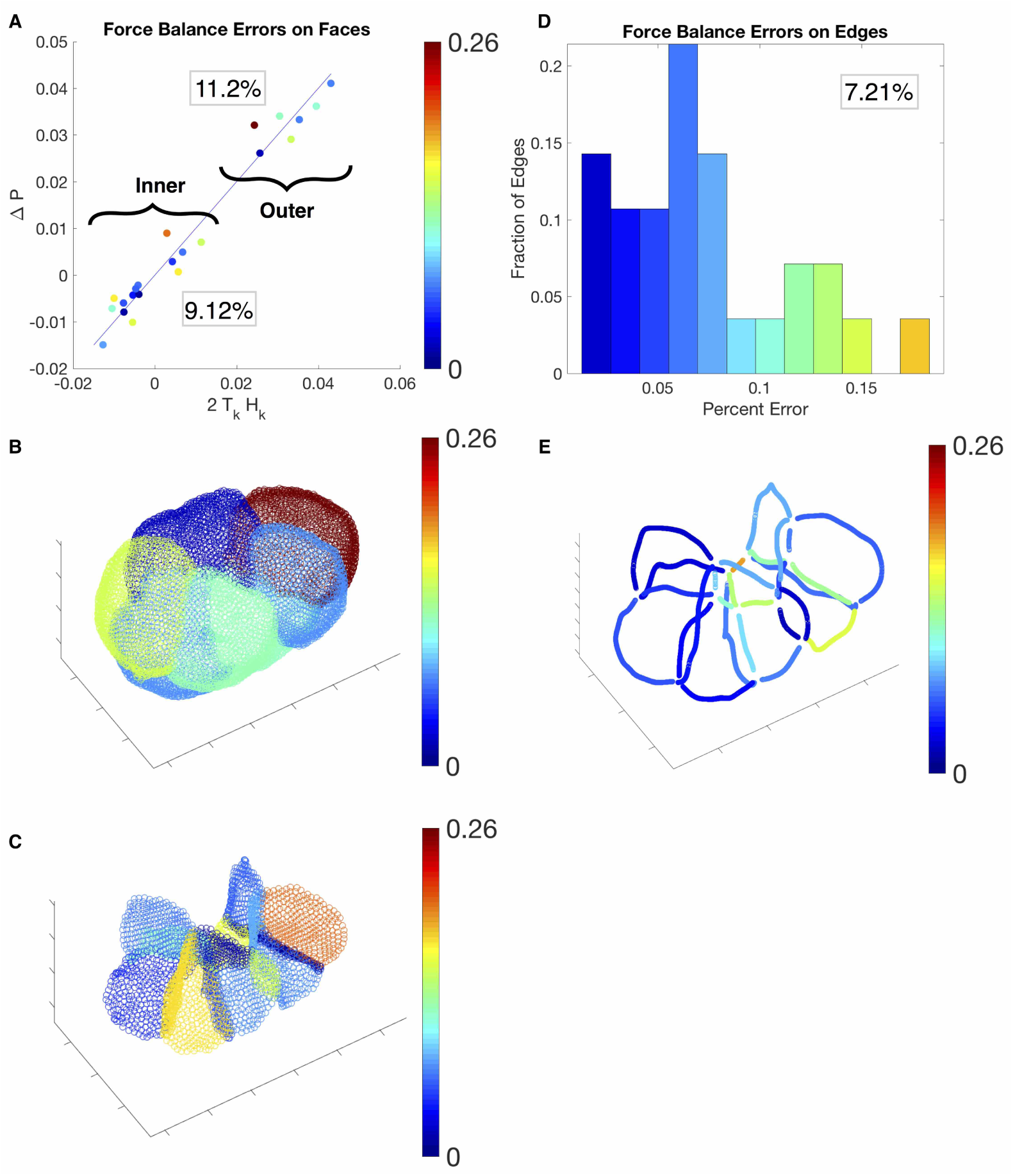
Error Plots for Inferred Forces. Plot of the force balance errors on the faces (A-C) and edge junctions (D-E). In A, we depict errors in Eq. (1) by plotting 2*T*_*k*_*H*_*k*_ against Δ*P* for each face in the scatter plot. The two clusters correspond to the inner and outer membranes, with their respective average percent errors in the box. Errors for the faces are portrayed as a heat map on the embryo, with outer membranes in B and inner membranes in C. In D, we plot the percent error equations (2) and (3) as a histogram, with the average percent errors in the box. The percent errors of edges are portrayed as a heat map on the edge junctions (E), where the heat map corresponds to the heat map in the histogram (D).

**Figure 10.**
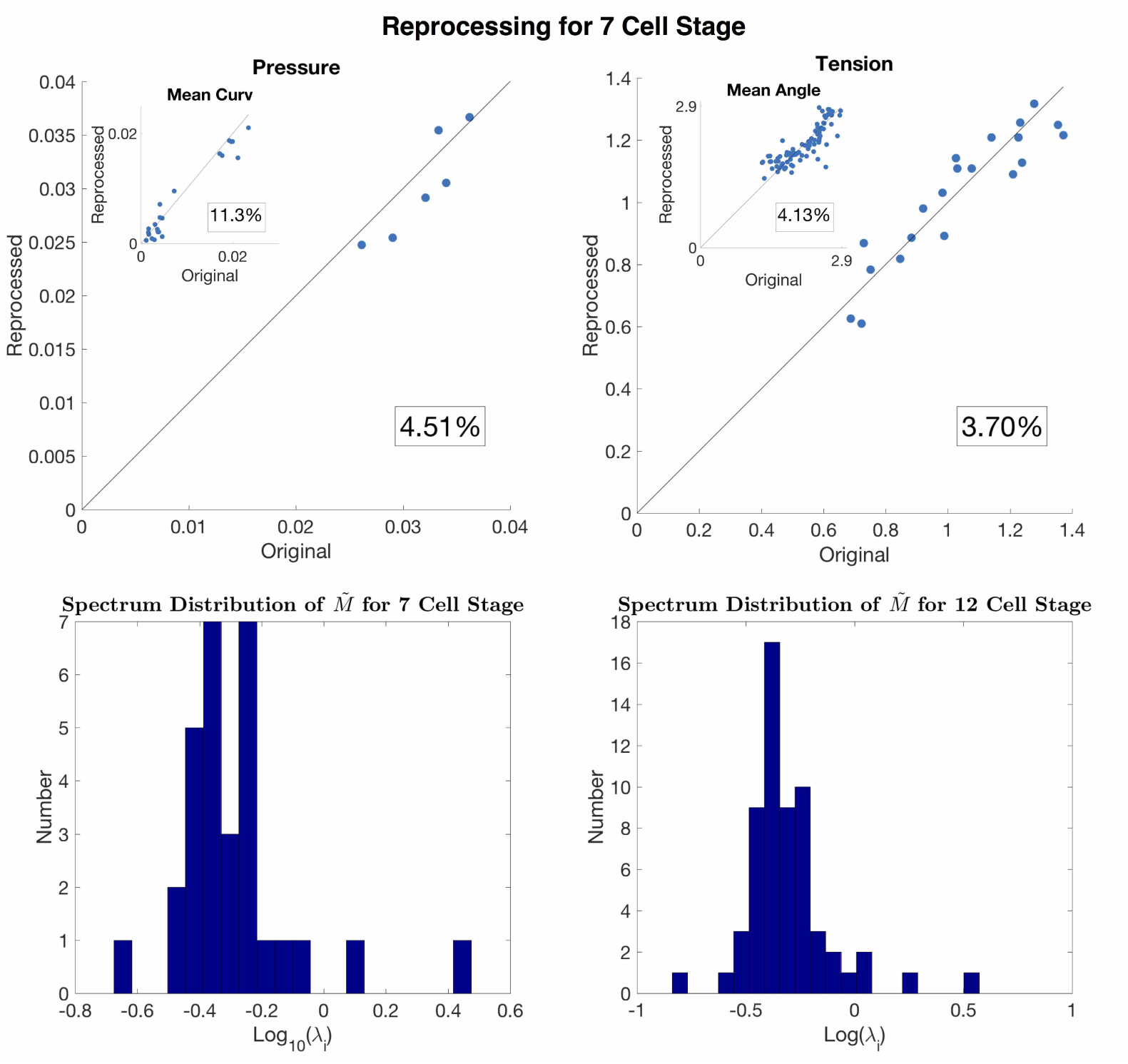
The sensitivity analysis and reproducibility. Plot of the error in the reprocessing at the 7 cell stage (top row). In the scatter plots, the original value are plotted against the reprocessed values, for the inferred pressure, tension, mean curvature, and mean angles. The percent error for each value is boxed. The eigenvalues of 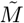 are plotted on a log scale histogram (bottom row) with the 7 cell stage on the left and 12 cell stage on the right.

## Discussion

In this study we have presented a morphological reconstruction scheme that forms the basis of a force inference method for analyzing the geometric and mechanical features of worm embryonic development. The novel morphological reconstruction scheme presented here has uses beyond facilitating force inference schemes. In particular, accurate membrane recognition permits quantitative measurements of signaling dynamics present at the membrane. Furthermore, the high resolution reconstruction of cellular geometries can form the basis of higher resolution force inference schemes that allow for inhomogeneities and anisotropies in membrane tensions. Additionally, our scheme facilitates the measurements of *in toto* velocity data that would be of interest to study during the processes of gastrulation and cell sorting in the worm. Furthermore, the pipeline developed here can be ported over for analysis of 3-dimensional live-imaging data more broadly. Foe example, the method could form the basis of an analysis of nuclear shape using a membrane or nuclear marker.

The rigorous assessment of the accuracy, reproducibility, and sensitivity of our force inference scheme highlights its strengths and weaknesses. The assessment also suggests improvements to the scheme that we are currently pursuing. More broadly speaking, we anticipate that force inference schemes will compliment the molecular tools under development to give us insight into morphogenesis. While FRET-based reporters, for example, give access to molecular level forces, connecting them to the processes of cell shape change and cell movement will require a model for how the two very different scales are connected. Force inference schemes on the other hand give insight into the forces that control gross cell shape and cell movement features but lack molecular insights. We anticipate that it will be combination of the aforementioned tools that will drive progress in the field.

## Supporting Information

### Some details in the reconstruction of the morphology

We start the morphological reconstruction by using standard machine learning image analysis software (*Ilastik*) to generate a probability map, evaluating the likelihood of a voxel point to be on the membrane or in the cytoplasm (or the perivitelline space exterior to the embryo), trained by grayscale images of the plasma membranes at different time points. We take the data into *MATLAB* and threshold the membrane probability map by *p*_*h*_ (see the parameter table for the values), considering values below *p*_*h*_ as 0. We then dilate the probability map on each voxel with a ball of radius *r*_*h*_, removing small regions on the membrane that is preconsidered as cytoplasmic regions by (*Ilastik*) followed by an erosion of the 0 *-* value voxel points with balls of radii *r*_*h*_. In addition, we have identified and removed connected membrane regions with size fewer than *V*_*min*_ voxels. We then perform a watershed transformation on the probability map to obtain a classification of the voxels into cells, separated by a one voxel thick representation of the membrane. The membrane can be segmented by dilating adjacent cells one voxel and retrieving their intersection. The edges can then be retrieved by dilating adjacent membrane faces one voxel and selecting the intersection. At last, the pipeline described above give rises to a data structure with cells, membrane faces and edge junctions and their connectivities.

### Some mathematical concepts relevant to the geometry of membranes

A membrane is topologically a surface embedded in the three-dimensional Euclidean space (**R**^3^). The mean curvature at a point *p* on the surface *S* is an invariant describing how the surface is bent in the **R**^3^. Here we summarize only relevant concepts from differential geometry, to clarify our procedure in computing the mean curvature in our work. Mean curvature is the mean between two principal normal curvatures. In the following, we explain the concept of curvature, the normal curvature, the principal curvatures and finally the mean curvature. We describe how to compute the mean curvature in the end.

### Curvature of a curve vs normal curvature of a surface

Given a curve **r**(*s*) embedded in **R**^3^ where *s* is the arc length along the curve, at a point *p* along the curve, the unit tangent vector is given as **r**^*′*^ (*s*) *|*_*s*=*p*_. Then **r**^*″*^ (*s*) is the rate of the change of unit tangent vector along *s*. We define the principal normal by

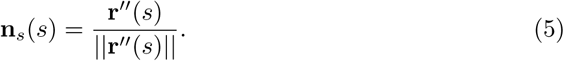

The curvature of **r**(*s*) is defined as

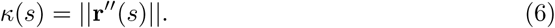

It is the rate of the change of tangent along the principal normal -**r**^*″*^ (*s*) = *κ*(*s*)**n**_*s*_(*s*). Now, given a curve **r**(*s*) = **r**(*u*(*s*)*, v*(*s*)) on a surface **r**(*u, v*), we can decompose **r**^*″*^ (*s*) by

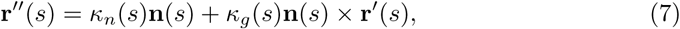

where **n**(*s*) is the surface normal, orthogonal to tangent vectors on the surface in all directions. *κ*_*n*_(*s*) is called the normal curvature and *κ*_*g*_(*s*) is called the geodesic curvature. What is interesting is that *κ*_*n*_(*s*) = **r**^*″*^ (*s*) *·* **n**(*s*) = *–***r**^*′*^ (*s*)*·* **n**^*′*^ (*s*), only depends **r**^*′*^ (*s*) and **n**^*′*^ (*s*), respectively the unit tangent vector and the rate of the change of the surface normal. It measures how is the surface bent, a property of the surface, instead of a curve on the surface. Notice that **n**^*′*^ (*s*) and **r**^*′*^ (*s*) are both in the tangent plane of the surface S at p, defined as *T*_*p*_*S*. The explanation is in the following subsection.

### The Weingarten map and the principal curvatures

The Weingarten map *W*_*p*_ is a unique linear map in *T*_*p*_*S* and can be determined by

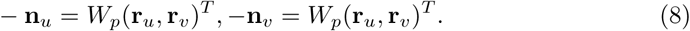

So we also realize −**n**^*′*^ (*s*) = *W*_*p*_**r**^*′*^ (*s*) and *?*_*n*_ = **r**^*′*^ (*s*) *· W*_*p*_**r**^*′*^*s*). Notice −**n**_*u*_ and −**n**_*v*_ are in *T*_*p*_*S* since −**n**_*u*_ *·* **n** = 0 and −**n**_*v*_ *·* **n** = 0. Given {**r**_*u*_, **r**_*v*_} as the basis, *W*_*p*_ is a 2 *×* 2 matrix. Since it is symmetric (shown later), there is always a pair of real eigenvalues *κ*_1_ and *κ*_2_ with the corresponding basis 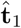 and 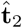 that satisfies

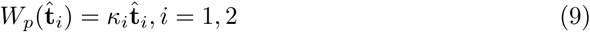

Notice *κ*_*i*_ is the normal curvature in the direction of 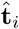. The pair of the normal curvatures are called the principal curvatures of the surface at *p*. The mean 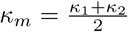 is mean curvature, and the product *κ*_*g*_ = *κ*_1_*κ*_2_ is the Gaussian curvature. They are both invariants of *W*_*p*_.

### The calculation of the mean curvature

We need to solve *W*_*p*_ and its eigenvalue pairs to compute the mean curvature 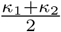. It can be solved by 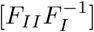 where

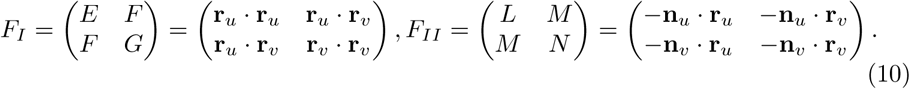

This can be shown by solving the linear map *W*_*p*_ determined by

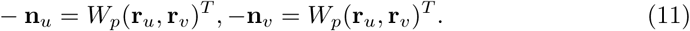

*F*_*I*_ and *F*_*II*_ are the matrix of the first and second fundamental form of the surface, respectively.

## Parameter Table

| Parameter | Value | Description |
| --- | --- | --- |
| $p_h$ | 0.5 | The membrane probability map threshold used in the reconstruction of the embryo. |
| $r_h$ | 3 (5) | The radius of the ball used in the morphological processes for the reconstruction of the embryo. |
| $V_{min}$ | 2000 (500) | The threshold for the minimum number of connected components used in the reconstruction of the embryo. |
| $K_S$ | 50 | The number of nearest neighboring points used for membrane smoothing. |
| $K_C$ | 50 (800, 3200) | The number of nearest neighboring points used for curvature computation on the membrane. |
| $d_T$ | 8 | The distance threshold in voxels from the other membranes from which a temporary junction point cloud is sampled. |
| $K_J$ | 240 | The number of nearest neighboring points used to thin the temporary junction point cloud. |

